# High-throughput amplicon sequencing of the full-length 16S rRNA gene with single-nucleotide resolution

**DOI:** 10.1101/392332

**Authors:** Benjamin J Callahan, Joan Wong, Cheryl Heiner, Steve Oh, Casey M Theriot, Ajay S Gulati, Sarah K McGill, Michael K Dougherty

## Abstract

Targeted PCR amplification and high-throughput sequencing (amplicon sequencing) of 16S rRNA gene fragments is widely used to profile microbial communities. New long-read sequencing technologies can sequence the entire 16S rRNA gene, but higher error rates have limited their attractiveness when accuracy is important. Here we present a high-throughput amplicon sequencing methodology based on PacBio circular consensus sequencing and the DADA2 sample inference method that measures the full-length 16S rRNA gene with single-nucleotide resolution and a near-zero error rate.

In two artificial communities of known composition, our method recovered the full complement of full-length 16S sequence variants from expected community members without residual errors. The measured abundances of intra-genomic sequence variants were in the integral ratios expected from the genuine allelic variants within a genome. The full-length 16S gene sequences recovered by our approach allowed *E. coli* strains to be correctly classified to the O157:H7 and K12 sub-species clades. In human fecal samples, our method showed strong technical replication and was able to recover the full complement of 16S rRNA alleles in several *E. coli* strains.

There are likely many applications beyond microbial profiling for which high-throughput amplicon sequencing of complete genes with single-nucleotide resolution will be of use.

## Introduction

The amplification of specific genetic loci by polymerase chain-reaction (PCR) can powerfully focus DNA sequencing on genetic variation of interest. Amplicon sequencing effectively detects genetic variation embedded in complex chemical and genetic backgrounds, and is far more cost-effective than untargeted sequencing when large amounts of undesired genetic material is present, as can be the case for host-associated microbial populations or specific genes in large genomes (Franzosa 2015). The precision, sensitivity and low cost of amplicon sequencing have made it a ubiquitous tool utilized in thousands of published scientific studies each year.

However, the advantages of amplicon sequencing come at a cost: the information provided by amplicon sequencing is limited to a single genetic locus. In current practice, the genetic loci measured by amplicon sequencing are typically restricted to 100–500 nucleotide regions that fit within the short reads generated by high-throughput sequencing platforms. In the popular community profiling application, short reads limit investigators to fragments of preferred taxonomic barcodes, such as the 16S rRNA gene in bacteria or the internal transcribed spacer (ITS) region in fungi, degrading taxonomic resolution and the ability to distinguish between related strains (Fuks 2018; Edgar 2018). In studies of functional genes, short reads do not cover even compact viral genes, limiting amplicon sequencing to incomplete measurements of functional genes.

In recent years, Pacific Biosciences (PacBio) and Oxford Nanopore have developed new technologies that generate long sequencing reads that can extend tens of thousands of nucleotides (Goodwin 2016; Levy 2016). Long reads can dramatically widen the genetic field of view measured by amplicon sequencing, offering the promise of greatly increased resolution in taxonomic profiling applications and measurement of complete functional genes. However, amplicon sequencing applications are often sensitive to the presence of spurious sequence variants introduced by PCR and sequencing errors, and long-read sequencing has a much higher error rate (~10%) than does short-read sequencing (~0.5%). For PacBio, high long-read error rates can be ameliorated by the construction of circular consensus sequences (CCS), in which an amplified genetic locus is circularized and read through multiple times before a consensus sequence is reported (Hebert 2018). The CCS approach effectively trades read length for accuracy: raw reads reaching over 100 kilobases in length are processed to yield CCS reads that extend only 1-20 kilobases but that have a per-base accuracy comparable to that of short-read sequencing (Jiao 2013; Larsen 2014; Wenger 2019).

Several previous studies have evaluated long-read amplicon sequencing of the 16S rRNA gene based on PacBio CCS and Oxford Nanopore reads, but have reported that a still considerable error rate necessitated binning similar sequences together to mitigate the impact of sequencing errors on subsequent analyses (Schloss 2016; Singer 2016; Wagner 2016; Schlaeppi 2016; Calus 2018). As a result, long-read amplicon sequencing was not found to improve resolution relative to short reads as much as was suggested by the increase in read length. The advantages of long reads have been a sufficient value proposition in applications such as tracking diversification of the HIV *env* gene (Caskey 2017; Eren 2018) and characterizing histocompatibility in non-human organisms (Westbrook 2015; Karl 2017), but the limited resolution gain in practice has been insufficient to justify the increased cost of long-read amplicon sequencing in many applications, and so it is not widely used at this time.

Here we introduce an amplicon sequencing methodology based on PacBio CCS sequencing and the DADA2 algorithm and software (Callahan 2016) that resolves exact amplicon sequence variants (ASVs) with single-nucleotide resolution (Callahan 2017) from the full-length 16S rRNA gene with a near-zero error rate. We evaluate the accuracy and precision of our method in mock microbial communities and in technically replicated samples from the human fecal microbiome. We demonstrate taxonomic classification at the sub-species level using the full complement of 16S rRNA gene alleles recovered by our method. Although our focus here is on the 16S rRNA gene, the core components of this methodology can be straightforwardly extended to other genetic loci for which suitable PCR primers are available.

## Materials and Methods

### Mock Communities

The “Zymo” mock community is the ZymoBIOMICS™ Microbial Community DNA Standard available from Zymo Research (Irvine, CA). The Zymo mock community consists of 8 phylogenetically distant bacterial strains (Table 1), 3 of which are gram-negative and 5 of which are gram-positive, as well as 2 yeast strains not amplified by our amplicon sequencing protocol. Genomic DNA from each bacterial strain is mixed in equimolar proportions. Of note, Zymo Research replaced five strains in the ZymoBIOMICS™ standards with similar strains beginning with Lot ZRC190633. The Lot # of the sample analyzed here was ZRC187325, which contains the old mixture of strains, including *E. coli* O157:H7 str. CDC B6914-MS1. Detailed information is available at the manufacturer’s website: https://www.zymoresearch.com/zymobiomics-community-standard.

**Table 1.** Bacterial strains included in the Zymo mock community. The species and 16S copy number of each of the eight bacterial strains included in the Zymo mock community are listed. Note that the two yeast strains included in the mock community were omitted, as yeasts are not amplified by our sequencing protocol. This information is adapted from Table 2 in the Technical Protocol for the ZymoBIOMICS™ Microbial Community DNA Standard (D6305).

| Species | GC Content (%) | 16S Copy # | Gram Stain |
| --- | --- | --- | --- |
| *Pseudomonas aeruginosa* | 66.2 | 4 | - |
| *Escherichia coli* | 46.7 | 7 | - |
| *Salmonella enterica* | 52.2 | 7 | - |
| *Lactobacillus fermentum* | 52.4 | 5 | + |
| *Enterococcus faecalis* | 37.5 | 4 | + |
| *Staphylococcus aureus* | 32.9 | 6 | + |
| *Listeria monocytogenes* | 38.0 | 6 | + |
| *Bacillus subtilis* | 43.9 | 10 | + |

**Table 2.** Bacterial strains included in the HMP mock community. The species and 16S copy number of the twenty bacterial strains included in the HMP mock community are listed. Species information is adapted from Table 2 in the Product Information Sheet for HM-783D from BEI resources.

| Species | 16S Copy # |
| --- | --- |
| *Acinetobacter baumannii* | 6 |
| *Actinomyces odontolyticus* | 2 |
| *Bacillus cereus* | 12 |
| *Bacteroides vulgatus* | 7 |
| *Clostridium beijerinckii* | 14 |
| *Deinococcus radiodurans* | 3 |
| *Enterococcus faecalis* | 4 |
| *Escherichia coli* | 7 |
| *Helicobacter pylori* | 2 |
| *Lactobacillus gasseri* | 6 |
| *Listeria monocytogenes* | 6 |
| *Neisseria meningitidis* | 4 |
| *Propionibacterium acnes* | 3 |
| *Pseudomonas aeruginosa* | 4 |
| *Rhodobacter sphaeroides* | 4 |
| *Staphylococcus aureus* | 6 |
| *Staphylococcus epidermidis* | 6 |
| *Streptococcus agalactiae* | 7 |
| *Streptococcus mutans* | 6 |
| *Streptococcus pneumoniae* | 4 |

The “HMP” mock community was obtained through BEI Resources, NIAID, NIH as part of the Human Microbiome Project: Genomic DNA from Microbial Mock Community B (Staggered, Low Concentration), v5.2L, for 16S rRNA Gene Sequencing, HM-783D. The HMP mock community consists of 20 bacterial strains (Table 2), most of which are phylogenetically distant from one other, but also contains two *Staphylococcus* species and three *Streptococcus* species. Genomic DNA from each bacterial strain was mixed so that the concentrations of rRNA gene operons vary over three orders of magnitude. Detailed information is available at the BEI website: https://www.beiresources.org/Catalog/otherProducts/HM-783D.aspx.

The 16S copy numbers of the strains in the HMP mock community were not provided by the vendor, so we determined 16S rRNA gene copy number of each strain by reference to the entries corresponding to that species in *rrn*DB version 5.4 (The Ribosomal RNA Database, https://rrndb.umms.med.umich.edu/, Stoddard 2015). Ambiguities were resolved by analyzing the number of 16S rRNA gene copies in the top BLAST hits of the ASVs assigned to each strain.

### DNA Extraction

The mock communities analyzed here consisted of genomic DNA extracted by the vendors. For the human fecal samples, we extracted genomic DNA with the MO Bio PowerFecal kit (Qiagen) automated for high throughput on QiaCube (Qiagen). The manufacturer’s instructions were followed with bead beating in 0.1 mm glass bead plates. DNA quantification was performed with the Qiant-iT Picogreen dsDNA Assay (Invitrogen).

### Amplicon Sequencing

The 27F:AGRGTTYGATYMTGGCTCAG and 1492R:RGYTACCTTGTTACGACTT universal primer set was used to amplify the full-length 16S rRNA gene from the genomic DNA extracted from each sample. For the fecal samples, both the forward and reverse 16S primers were tailed with sample-specific PacBio barcode sequences to allow for multiplexed sequencing. The fecal samples were barcoded to allow multiplexed sequencing. We chose to use barcoded primers because this reduces chimera formation as compared to the alternative protocol in which primers are added in a second PCR reaction. The KAPA HiFi Hot Start DNA Polymerase (KAPA Biosystems) was used to perform twenty cycles of PCR amplification, with denaturing at 95C for 30 seconds, annealing at 57C for 30 seconds, and extension at 72C for 60 seconds. Post-amplification quality control was performed by on a Bioanalyzer (Agilent Technologies, Santa Clara, CA). Amplified DNA from the fecal samples was then pooled in equimolar concentration.

SMRTbell libraries were prepared from the amplified DNA by blunt-ligation according to the manufacturer’s instructions (Pacific Biosciences). Purified SMRTbell libraries from the Zymo and HMP mock communities were sequenced on dedicated PacBio Sequel cells using the S/P1-C1.2 sequencing chemistry. Purified SMRTbell libraries from the pooled and barcoded fecal samples were sequenced on a single PacBio Sequel cell. Replicate 1 of the fecal samples was sequenced using the S/P2-C2/5.0 sequencing chemistry, and Replicate 2 of the fecal samples was sequenced with a pre-release version of the S/P3-C3/5.0 sequencing chemistry.

Full details are presented in the Full-Length 16S Amplification SMRTbell® Library Preparation and Sequencing Procedure & Checklist included as Supplementary Document 1. For investigators interested in applying our methodology to other genes, we have included the more general-purpose Preparing Amplicon Libraries using PacBio^®^ Barcoded Adapters for Multiplex SMRT^®^ Sequencing Procedure & Checklist that is suitable for use with any amplicon as Supplementary Document 2. All amplicon sequencing was performed by Pacific Biosciences Inc. (Menlo Park, CA).

### Defining and Curating CCS reads

Circular consensus sequence (CCS) reads were generated from the raw PacBio sequencing data using the standard software tools provided by the manufacturer (Pacific Biosciences). CCS reads from the Zymo and HMP mock communities were generated using the *ccs* application with *minPasses*=*3* and *minPredictedAccuracy*=*0.999* in the SMRT Link 3.1.1 software package (Pacific Biosciences, 2016). For the fecal samples, raw reads were first demultiplexed using the *lima* application, specifying that the same barcodes were attached at both ends of an insert using the flags *same* and *peek-guess*, followed by generation of CCS reads using the *ccs* application with *minPasses*=*3* and *minPredictedAccuracy*=*0.999* in the SMRT Link 5.1 software package (Pacific Biosciences, 2018). Of note, the *ccs* application was updated in SMRTLink 3.0 to use the superior Arrow model (Hepler 2016), and CCS reads generated by the earlier SMRT Portal software may not be as accurate.

### Algorithms and Software

The DADA2 method is an algorithm for the inference of the ASVs (amplicon sequence variants, i.e. the true error-free sequences) present in a sample from the library of noisy reads generated by amplicon sequencing. DADA2 was initially developed for short-read amplicon sequencing, and a detailed description of the algorithm is available in the original publications (Rosen 2012; Callahan 2016). Briefly, the DADA2 algorithm uses a statistical model of the amplicon sequencing error process to identify sequences that are repeatedly observed too many times to be consistent with being generated by amplicon sequencing errors, and thus must represent distinct true sequences present in the sample. The DADA2 error model incorporates the quality information associated with each read, and thus repeated observations of sequences that are differentiated from others only at low-quality base positions are discounted appropriately. The key assumptions made by the DADA2 error model are that the relationship between the quality score and the likelihood of an error is consistent across all base positions in a read, and that after accounting for correlations in quality scores and nucleotide composition that errors are independent between reads.

The DADA2 R software package implements the DADA2 algorithm as part of a complete workflow that takes raw amplicon sequencing data in fastq files as input, and produces an error-corrected table of the abundances of amplicon sequence variants (ASVs) in each sample as output (Figure 1; Callahan 2016; Callahan 2017). The standard processing steps in the DADA2 workflow include quality filtering, dereplication, learning the dataset-specific error model, ASV inference, chimera removal, and taxonomic assignment.

**Figure 1:**
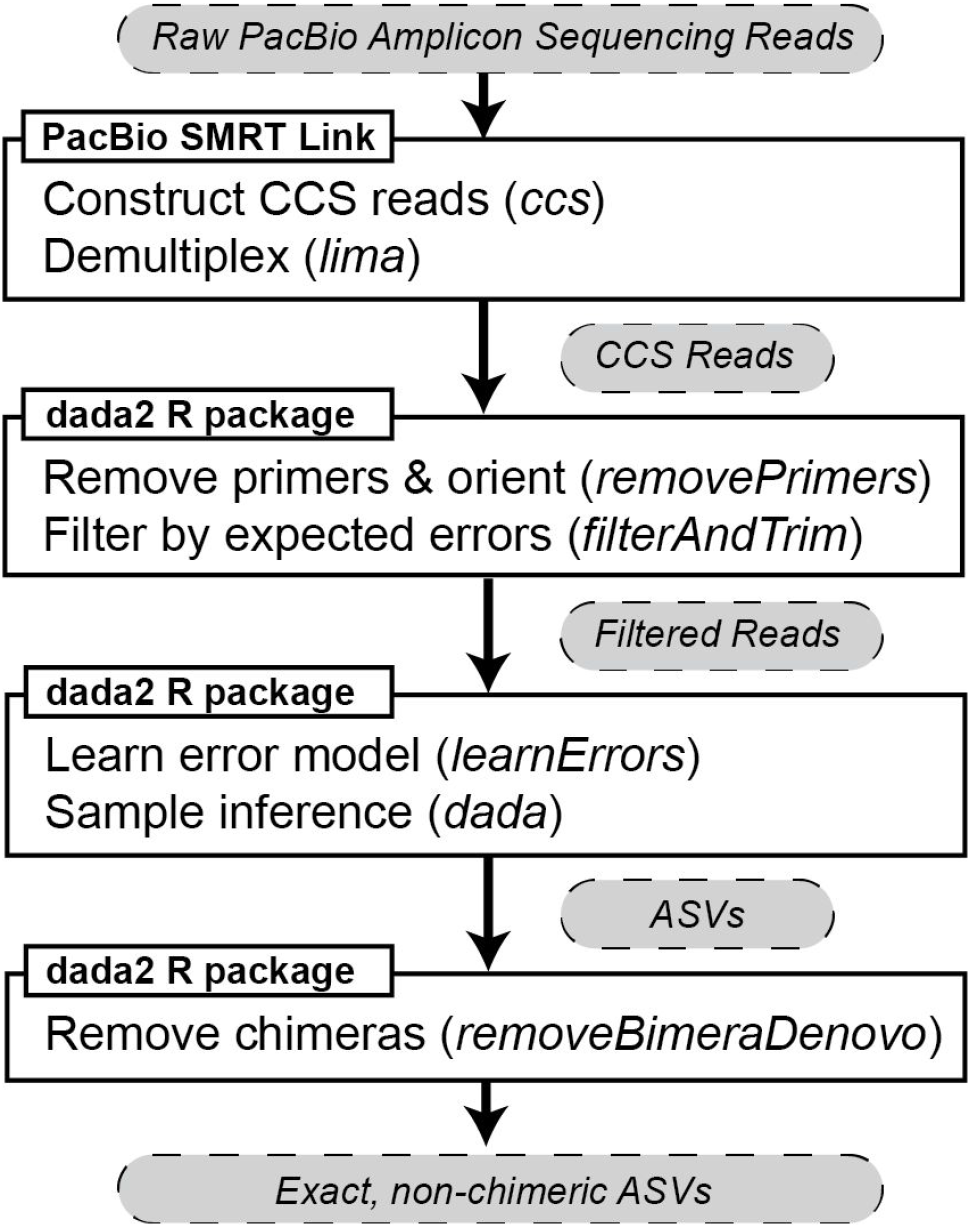
Flowchart of computational methodology. Two software packages, PacBio SMRT Link and the DADA2 R package, were used to process raw PacBio amplicon sequencing data into chimera-free amplicon sequence variants (ASVs). The specific commands used at each step of data processing are italicized, and the data types at each stage of processing are indicated in grey bubbles. Complete examples of this workflow are available in the reproducible analyses accompanying this manuscript: https://github.com/benjjneb/LRASManuscript.

The DADA2 R software package was improved and extended in a number of ways to enable precise ASV inference from long amplicon reads. The core data structures and sequence comparison algorithms used by the DADA2 algorithm were augmented to handle quality scores up to 93, sequence lengths up to 3000 nucleotides, and variable sequence lengths. A new error estimation procedure was introduced for PacBio CCS data, in which estimation of the error rate for bases with the maximum quality score of 93 is separated from estimation of error rates over the remainder of the quality score distribution. A new *removePrimers* function was added to the DADA2 R package that detects and removes primers from PacBio CCS reads, and orients them in the forward direction, as CCS reads are generated in a mixture of forward and reverse-complement orientations. A new default value of *minFoldParentOverabundance*=*3*.5 is recommended for chimera identification in full-length 16S rRNA gene sequencing data, in order to avoid spurious identification of some 16S variants as chimeras of other 16S variants that occur at higher copy number within the same genome.

The DADA2 R software package was improved and extended in a number of ways to enable efficient processing of large long-read amplicon sequencing datasets. Multithreading has been implemented for all computationally expensive steps in the DADA2 workflow, including filtering, error-model learning, ASV inference, chimera removal and taxonomic assignment. The *multithread* parameter in these functions controls the number of threads, and *multithread*=*TRUE* will automatically detect and use all available threads. Pairwise sequence comparison has been accelerated in several ways. The initial calculation of kmer-distances from the overlap between lexically-ordered kmer arrays has been explicitly vectorized using SSE2 instructions, achieving close to the theoretically maximum 16X speedup. If the lexically-ordered kmer arrays were sufficiently similar, a new step has been added in which the overlap between sequence-ordered kmer arrays is calculated using explicitly vectorized SSE2 instructions. If the overlap between the lexically-ordered kmer arrays is equal to the overlap between the sequence-ordered kmer arrays then the ungapped alignment is assumed to be the optimal alignment between the sequences. Otherwise a banded Needleman-Wunsch alignment is performed that has been implicitly vectorized along the sub-diagonal.

### Code Availabilty

The DADA2 R software package is licensed under the LGPL3, and the source code is freely available from GitHub: https://github.com/benjjneb/dada2. Reproducible R markdown documents implementing the analyses performed in this paper also document the full workflow used to infer ASVs from PacBio CCS data: https://github.com/benjjneb/LRASManuscript. All computation was performed with version 1.12.1 of the DADA2 R software package on a 2017 MacBook Pro and was completed in minutes to tens of minutes.

### Full-Complement Classification of *E. coli*

ASVs assigned to the *Escherichia* genus were grouped into strain-level bins based on the expectation of integral ratios between same-genome ASVs and the known copy number of seven 16S rRNA genes in *E. coli*. We performed BLAST searches for each ASV in a strain-level bin against the NCBI nucleotide database (nt) on February 4, 2019. The highest BLAST score for each ASV was determined, and all accessions reaching the highest score were recorded. The total occurrences of accessions across the high scores for each ASV were tabulated. The set of accessions with the greatest number of high-score hits served as the basis for further classification. The metadata associated with the Genbank entry for each such accession was inspected, and *E. coli* strains were assigned metadata values (e.g. serotype) that were consistent across all such accessions. Accessions for which the metadata entry of interest was absent were ignored.

Classification based on representative full-length sequences pursued the same BLAST-and-consensus approach as above, but utilized only the most abundant ASV from each strain as the representative sequence. Classification based on the V3V4 and V4V5 gene regions followed the same procedure as full-complement classification, but using the subsequences extracted using the Biostrings *matchLRPatterns* function (Pagès 2017) based on the Illumina-recommended V3V4 primer set and the JGI-recommended V4V5 primer set, respectively (Klindworth 2013; Parada 2016).

### Determination of Genomic Abundances

We used the 16S rRNA gene copy number, along with the frequencies of the associated ASVs, to infer the genomic frequency of each mock community strain in the pool of amplified DNA. More precisely, to calculate the genomic frequencies we summed the frequencies of all ASVs assigned to each strain, and then divided by the corresponding 16S rRNA gene copy number.

### Determination of Sequence Accuracy

The available reference databases for the Zymo and HMP mock communities are incomplete, so we used a multifaceted approach to evaluate the accuracy of denoised ASVs. First, ASVs were compared to the incomplete reference databases associated with each mock community. Second, each ASVs was BLAST-ed against nt excluding uncultured/environmental sample sequences. Finally, the genomic abundance of each ASV was calculated (see above). An ASV was determined to be accurate if it exactly matched a sequence previously observed from an isolate of the corresponding species, or if the ASV appeared in an integer number (±0.2) of copies per genome and there were multiple alleles present in the data from that strain.

## Results

### Accuracy in Mock Communities of Known Composition

We evaluated the performance of our full-length 16S rRNA amplicon sequencing methodology in two artificially constructed communities of known composition (mock communities). The Zymo mock community contains eight phylogenetically distant bacterial strains, with cellular proportions chosen to equalize the total genomic DNA contributed by each strain. The HMP mock community contains 20 bacterial strains of varying phylogenetic similarity, with proportions chosen such that rRNA gene frequencies vary over 3 orders of magnitude. Each mock community was sequenced on a single Sequel cell using the S/P1-C1.2 sequencing chemistry. They Zymo mock community yielded 77,453 CCS reads above the *minPasses* threshold of 3 and the *minPredictedAccuracy* threshold of 99.9%, and 69,367 reads after removing primers and filtering (Methods, Supplementary Table 1). The HMP mock community yielded 78,328 CCS reads above the thresholds, and 69,963 reads after removing primers and filtering.

Exact amplicon sequence variants (ASVs) were inferred from the filtered reads by the new version 1.12.1 of the DADA2 R software package that has been updated to efficiently process long amplicon reads and appropriately model PacBio CCS sequencing errors (Methods). Taxonomy down to the genus level was assigned to ASVs by the naive Bayesian classifier and the SILVA v128 database (Wang 2007; Quast 2013), and at the species level by BLAST searches against the NCBI nucleotide collection (nt). In both datasets, every ASV was assigned to a genus and species belonging to the expected members of the mock communities. Since we expect to detect intragenomic variation in the often multi-copy 16S rRNA gene, we grouped ASVs into putative strain bins by their taxonomic assignments.

In the Zymo mock community 29 ASVs were detected, of which 25 of 29 were exact matches (100% identity, 100% coverage) to 16S rRNA genes previously sequenced from the isolates of the corresponding species. The three *Lactobacillus fermentum* ASVs and one *Staphylococcus aureus* ASV that were not exact matches differed by just one nucleotide from previously observed sequences. In the HMP mock community 51 ASVs were detected, of which 48 of 51 were exact matches. The one *Staphylococcus epidermis* ASV and two *Clostridium bejerenckii* ASVs that were not exact matches differed by just two and one nucleotides, respectively, from previously observed sequences. The frequencies of ASVs detected in the HMP mock community varied over 3 orders of magnitude, ranging as low as 0.00019.

If ASVs represent genuine allelic variants present in these genomes, then ASVs from the same strain should appear in integral ratios corresponding to the copy number of each allele within the genome. Integral ratios are unexpected if ASVs represent uncorrected amplicon sequencing artefacts. The ASVs inferred by our methodology appear in frequencies that are integral ratios to one another, and to the genomic frequency of the corresponding strain. Figure 2 shows the frequency of each ASV in the Zymo mock community, normalized to the observed strain genomic frequency (Methods). All ASV:genome ratios are integers (e.g. 1, 2, 3, …) with maximum deviations of ±0.2. The frequencies of ASVs in the HMP mock community also appear in integral ratios, with the exception of a deviation in *Bacillus cereus* ASV frequencies that cannot be attributed to counting noise, given the substantial abundance of that strain in the data (Figure 3).

**Figure 2.**
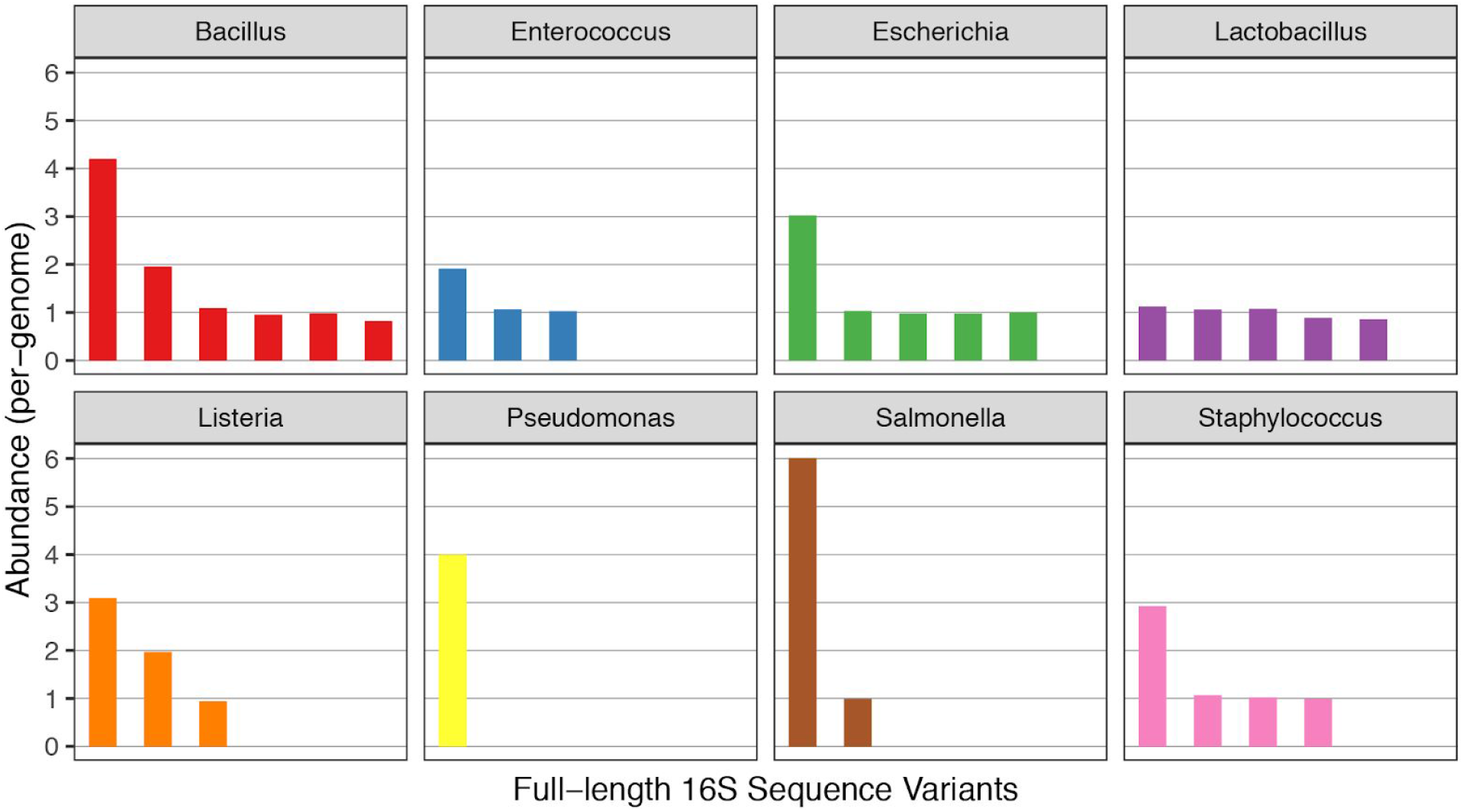
Abundances of full-length 16S rRNA gene amplicon sequence variants (ASVs) detected in the Zymo mock community, scaled by the genomic abundance. Twenty-nine distinct ASVs were detected by our long-read amplicon sequencing methodology in the Zymo mock community. Each ASV was grouped into a bin corresponding to eight bacterial strains in the mock community on the basis of its taxonomic assignment. The abundance of each ASV was divided by the genomic abundance of the mock community strain from which it originated (see Methods), and the normalized abundance of each ASV is plotted on the y-axis. Panels correspond to the 8 bacterial strains in the Zymo mock community, most of which were associated with multiple unique ASVs, i.e. distinct alleles from different copies of the rrn operon. No other ASVs were detected.

**Figure 3.**
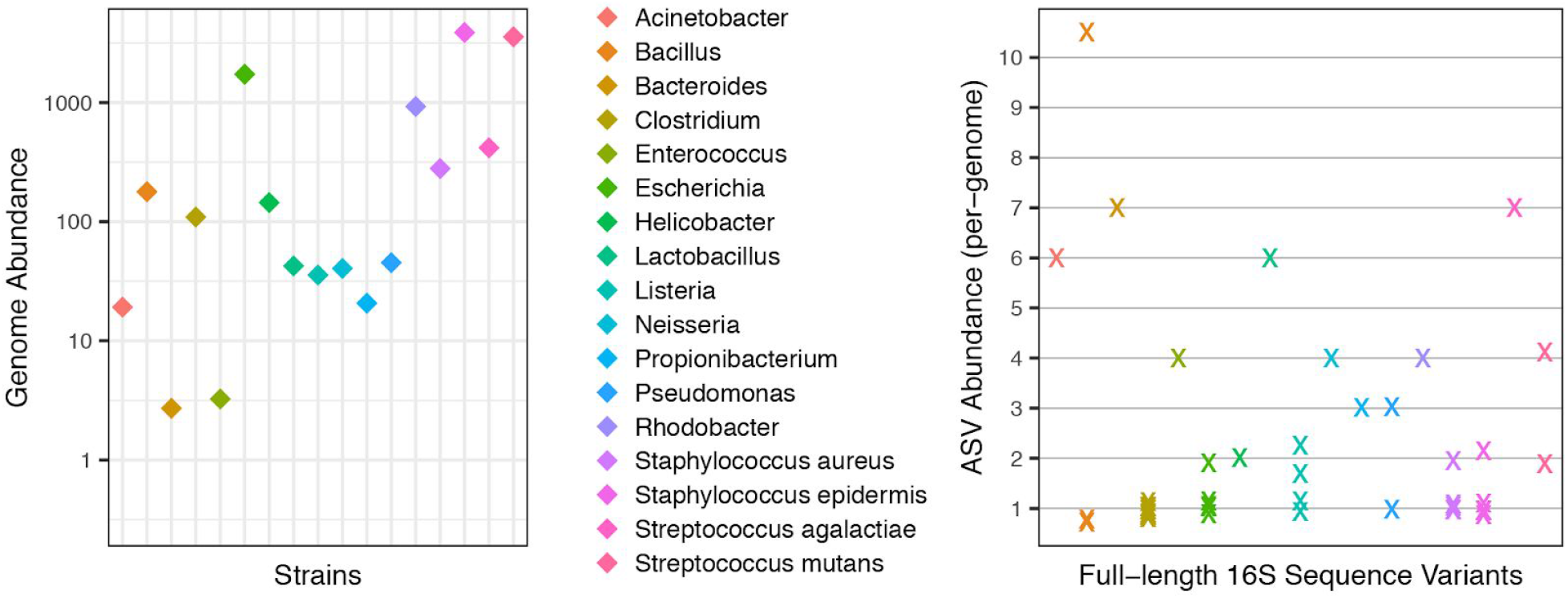
Abundances of genomes and ASVs recovered from the HMP mock community. (a) The abundances of each genome in the long-read amplicon sequencing data are plotted on the log-scaled y-axis (Methods). Observed genome abundances varied over three orders of magnitude. Significant counting noise exists for ASVs from genomes with abundances below 100. (b) The abundance of each ASV divided by the genomic abundance of the mock community strain from which it originated is plotted on the y-axis. Integral values are indicated by horizontal grid lines. No other ASVs were detected.

### Error Rates of PacBio CCS Amplicon Reads and Denoised ASVs

The available reference databases for these mock communities are incomplete, so we used a multifaceted approach to evaluate ASV accuracy (Methods) that incorporated both comparisons to reference databases and the pattern of integral ratios expected between genuine allelic variants within a genome. In the Zymo mock community, we concluded that the ASVs detected by our methods contained no false positives and that the residual per-base error rate after denoising with DADA2 was zero. In the HMP mock community, we concluded that it was most likely that the ASVs detected by our method contained no false positives, in which case the residual per-base error rate of the denoised ASVs was again zero. However, we leave open the possibility that the less frequent *B. cereus* ASVs that did not appear in integral ratios could contain errors, in which case the residual per-base error rate would be 2.6 × 10^−6^.

The Zymo mock community was used to characterize in detail the error profile of the uncorrected PacBio CCS amplicon reads generated by our protocol. After excluding chimeras and contaminants, the remaining reads were aligned to the true ASVs from which they were most likely to have originated. All differences between reads and true sequences were recorded by type (i.e. substitution, insertion or deletion), position in the read, and associated quality score. The measured error rates are shown in Figure 4, restricted to a representative window of 200 nucleotides for visual clarity. A complete plot over the full 16S rRNA gene amplicon in a vector image format suitable for zooming is included as Supplementary Figure 1.

**Figure 4.**
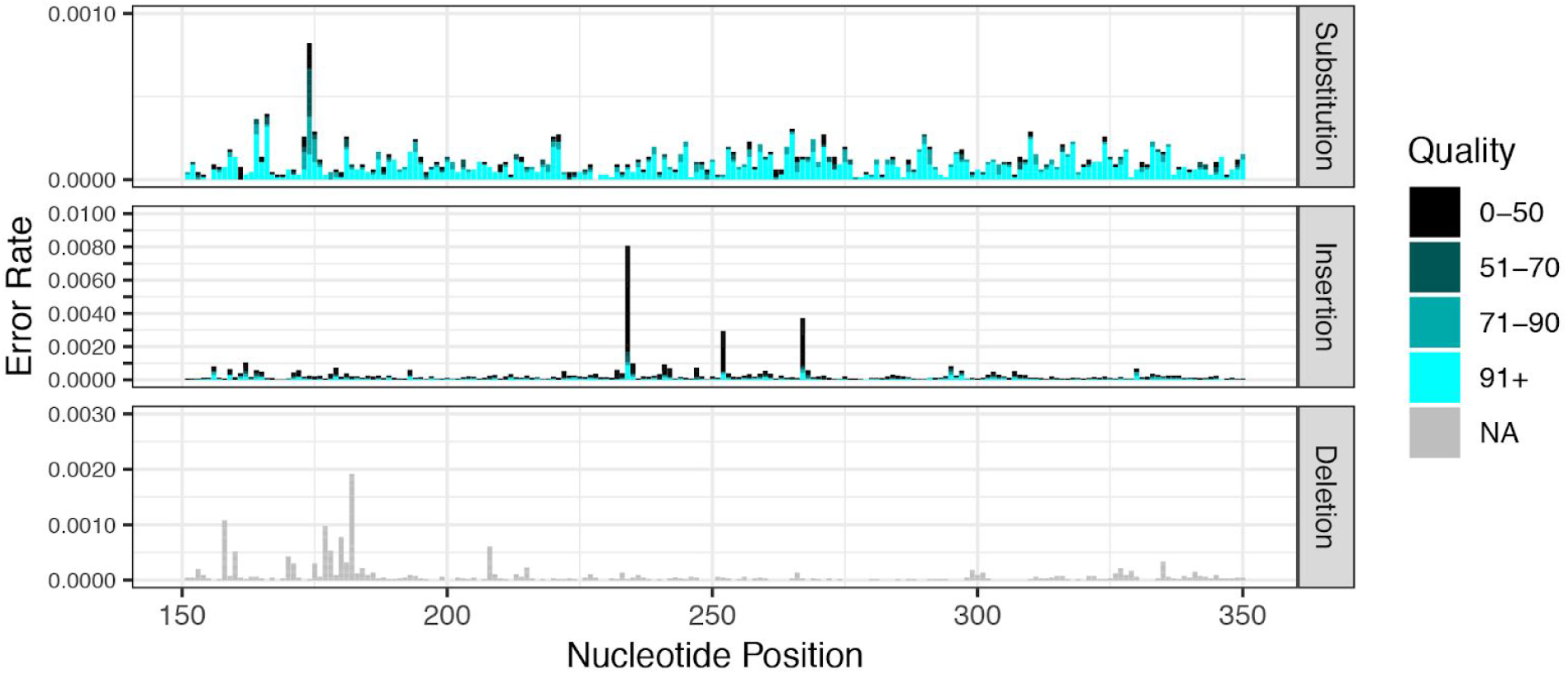
Error rates in PacBio CCS amplicon reads from the Zymo mock community as a function of error type, nucleotide position and quality score. The rate of substitutions (top), insertions (middle), and deletions (bottom) is shown for non-chimeric and non-contaminant reads from the Zymo mock community. Darker colors indicate lower quality bases. There is no quality score associated with deletions, as such errors indicate the absence of a nucleotide in the sequencing read. For visual clarity, only nucleotide positions 151-350 are plotted. See Supplementary Figure 1 for the full-length plot.

The aggregate error rate of the uncorrected CCS reads was 4.3 × 10^−4^ per nucleotide, with insertions (2.2 × 10^−4^) occurring at about twice about the rate of substitutions (1.1 × 10^−4^) and deletions (1.0 × 10^−4^). The distribution of substitutions across nucleotide positions within reads was relatively uniform, with a very small number of higher error rate positions that were associated with lower quality scores (Figure 4 and Supplementary Figure 1). The rate of insertion errors varied more across positions, likely due to context-dependent error modes such as homopolymer errors, but high error-rate positions were strongly associated with low quality scores. The variation in the rate of deletions across positions was intermediate between substitutions and insertions and, because deletions have no associated quality score, the high rates of deletions at certain positions was not obviously explained by a technical variable.

Earlier evaluations of full-length 16S rRNA gene amplicon sequencing using PacBio CCS reads generated by the RSII sequencing instrument reported far higher error rates before and after denoising, and perhaps more importantly also reported the presence of systematic errors repeatedly arose at specific positions (Schloss 2016; Wagner 2016). We re-analyzed the sequencing data from the *Staphylococcus aureus* monoculture data used to describe systematic errors in Wagner et al. 2016 with the updated DADA2 R software package, and recovered 5 non-chimeric ASVs. The *S. aureus* genome typically contains 5 copies of the *rrn* operon, and all 5 ASVs exactly matched previously sequenced *S. aureus* 16S rRNA genes. The differences between these intragenomic variants may have been misinterpreted as systematic errors, perhaps because the short-read genome assembly that was used as the ground truth in the original publication contained only one of the five *rrn* operons in the *S. aureus* genome (Larner-Svensson 2013). Additionally, we did not see an improvement in the accuracy of ASVs recovered from our mock communities when requiring higher subread coverage than the default threshold of *minPasses*=*3*, suggesting that the default thresholds are effective for Sequel sequencing chemistries and current software versions.

### Comparison of the Accuracy of DADA2 ASVs to Long-Read OTU Methods

We compared the accuracy of our DADA2-based computational pipeline to two other pipelines that have previously been applied to PacBio amplicon sequencing data, one based on the mothur software package (Schloss 2009; Schloss 2016) and one based on the uparse algorithm implemented in the usearch software package (Edgar 2013; Earl 2018). Each computational pipeline was applied to commonly pre-processed fastq files from the Zymo and HMP mock community datasets. The accuracy of the ASVs output by DADA2 and the OTUs output by mothur and uparse are reported in Table 3.

**Table 3.** The accuracy of different computational methods on PacBio amplicon sequencing data from mock communities. The PacBio long-read amplicon sequencing data from the Zymo and HMP mock communities was processed by DADA2, mothur and uparse (as implemented in usearch). DADA2 identifies exact amplicon sequence variants (ASVs) with single-nucleotide resolution, while mothur and usearch identify OTUs, i.e. clusters of reads within a 97% similarity threshold. Total: The number of features (ASVs or OTUs) identified. TP: True positives. FP: False positives. Singleton: Features with just one read. In these datasets, the singleton OTUs output by mothur consisted of chimeras, sequencing errors and contaminants. Strains detected: The number of strains in the mock communities identified by each method.

| Dataset | Method | Feature Type | Total | TP | FP | Singleton | Strains Detected |
| --- | --- | --- | --- | --- | --- | --- | --- |
| Zymo | DADA2 | ASVs | 29 | 29 | 0 | 0 | 8 |
|  | mothur | OTUs | 24 | 8 | 5 | 11 | 8 |
|  | uparse | OTUs | 8 | 8 | 0 | 0 | 8 |
| HMP | DADA2 | ASVs | 51 | 51 | 0 | 0 | 17 |
|  | mothur | OTUs | 29 | 16 | 4 | 9 | 16 |
|  | uparse | OTUs | 16 | 16 | 0 | 0 | 16 |

**Table 4.** Classification of *E. coli* strains based on full-length and short-read 16S rRNA gene amplicon sequence variants. The ASVs identified from the E. coli strains in the Zymo and HMP mock communities were used to classify the strains based on the V3V4 region alone, the V4V5 region alone, the representative (i.e. most abundant) full-length (FL) sequence, and the full complement of full-length sequences (Methods). Classification was based on a consensus across the best hits identified by exact BLAST hits against the NCBI nucleotide database (nt). The number of equally best hits is recorded.

|  | Method | Classification | Best Hits |
| --- | --- | --- | --- |
| Zymo | V3V4 | *Escherichia/Shigella* | 994 |
|  | V4V5 | *Escherichia/Shigella* | 996 |
|  | FL-Representative | *E. coli* O157:H7 | 53 |
|  | FL-Full Complement | *E. coli* O157:H7 | 14 |
| HMP | V3V4 | *Escherichia/Shigella* | 994 |
|  | V4V5 | *Escherichia/Shigella* | 996 |
|  | FL-Representative | *E. coli* | 320 |
|  | FL-Full Complement | *E. coli* K12 | 32 |

All computational methods performed reasonably well, reflecting the high accuracy of the underlying PacBio CCS reads, but clear differences between the methods were also apparent. Both DADA2 and uparse had perfect specificity, in that every ASV or OTU output by these methods was truly present in the expected mock communities. The mothur method output 16 and 13 false positive OTUs in the Zymo and HMP mock communities, respectively. Most of these false positives were present as singletons (i.e. OTUs with just one read) and therefore would be removed if a singleton filter were applied prior to subsequent analysis. All three methods detected all 8 strains in the Zymo mock community. DADA2 detected 17 strains in the HMP mock community while mothur and uparse detected only 16 strains. DADA2 was able to distinguish between between *Staphylococcus aureus* and *Staphylococcus epidermidis* while mothur and uparse lumped these species together into a single OTU. Mothur also failed to detect the moderately abundant (>100 reads) *Neisseria gonorrhoeae* strain, but was the only method that detected the rare (3 reads) *Deinococcus radiodurans* strain.

### Sub-species Classification using the Full Complement of 16S rRNA Gene Alleles

Taxonomic classification is typically performed on 16S rRNA sequences individually, but greater resolution can be achieved by utilizing the full complement of 16S alleles in bacterial strains. In the Zymo mock community, 5 *E. coli* ASVs were recovered with abundances in 3:1:1:1:1 ratios. In the HMP mock community, 6 *E. coli* ASVs were recovered with abundances in 2:1:1:1:1:1 ratios. We consider here a simple *ad hoc* full-complement classification procedure in which the accessions of the best BLAST hits for each *E. coli* ASV against the NCBI nucleotide database (nt) are recorded, and sub-species classifications are assigned based on the set of accessions that were best hits to the largest number of ASVs (Methods). We compared full-complement classification to classification based on the most abundant (i.e. representative) sequence from each strain, as well as classification based on short-read ASVs from the commonly used V3V4 and V4V5 regions of the 16S rRNA gene (Table 3).

The exact full-length 16S rRNA gene sequences identified by our method allowed for the *E. coli* strain in the Zymo mock community to be classified as belonging to the enterohemorrhagic O157:H7 clade. Classification precision increased further when making use of the full complement of 16S rRNA alleles. There were 14 accessions that exactly matched 4 of the 5 ASVs recovered from the Zymo *E. coli* strain, and none that exactly matched all five. Of those 14 accessions, 12 were annotated with serotype O157:H7, one was annotated with O antigen 157 but had no annotation for the H antigen, and one had no serotype annotation. If we ignore the missing values, the Zymo *E. coli* strain was unambiguously and correctly classified as belonging to the enterohemorrhagic O157:H7 clade. In comparison, the short-read V3V4 and V4V5 amplicon gene sequences did not allow resolution between the *Escherichia* and *Shigella* genera, much less sub-species classification.

The full complement of full-length 16S rRNA gene sequences identified by our method allowed the *E. coli* strain in the HMP mock community to be classified as belonging to the K12 clade. There were 32 accessions that exactly matched all 6 ASVs recovered from the HMP *E. coli* strain. Of those 32 accessions, 27 were either annotated as K-12, MG1655 (a specific strain in the K-12 clade), or as derived from MG1655. The other 5 were unannotated, but the best BLAST hits to the full genomes were K-12 strains. Classification based on only the most abundant full-length sequence did not resolve this clade membership, and allowed only classification to the *E. coli* species. The short-read V3V4 and V4V5 sequences of this K12 strain were identical to those obtained from the Zymo O157:H7 strain, and once again did not allow resolution between the *Escherichia* and *Shigella* genera.

### Resolution and Technical Replication in Human Fecal Samples

To validate the performance of our methodology in complex microbial communities, we performed technical replicate long-read amplicon sequencing experiments on a set of 9 human fecal samples and 3 additional samples excluded from further analysis. Individual samples were barcoded, and multiplexed sequencing of the 12 samples was performed in a single Sequel cell. The first replicate was sequenced using the S/P2-C2/5.0 Sequel sequencing chemistry, which yielded 177,691 CCS reads at 99.9% predicted accuracy across the twelve samples, 146,589 reads after primer detection and filtering, and a median of 12,158 filtered reads per human fecal sample (Supplementary Table 1). The second replicate was sequenced using a pre-release version of S/P3-C3/5.0 Sequel sequencing chemistry, which yielded 289,644 CCS reads at 99.9% predicted accuracy across the twelve samples, 249,802 reads after primer detection and filtering, and a median of 21,411 filtered reads per human fecal sample, nearly double that achieved using the previous generation sequencing chemistry.

We do not know the ground truth of these human fecal samples, so as an alternative to accuracy analysis we investigated consistency across technical replicates. Prior to sample inference with DADA2, we rarefied each replicate sample to 10,000 reads in order to remove any impact of sequencing depth, yielding 7 pairs of samples in which both replicates exceeded 10,000 reads. The ASVs detected by DADA2 from these rarefied samples were consistent across technical replicates (Figure 5A). On a per-sample basis, 881 ASVs were detected in both replicates, while 65 and 70 were detected only in Replicate 1 or 2, respectively. Estimated abundances were highly consistent: the Pearson’s correlation between the per-sample abundances across replicates was 0.998. ASVs that failed to be detected in one replicate or the other appeared at low frequencies (<50 reads, <0.5% frequency; Figure 5A).

**Figure 5.**
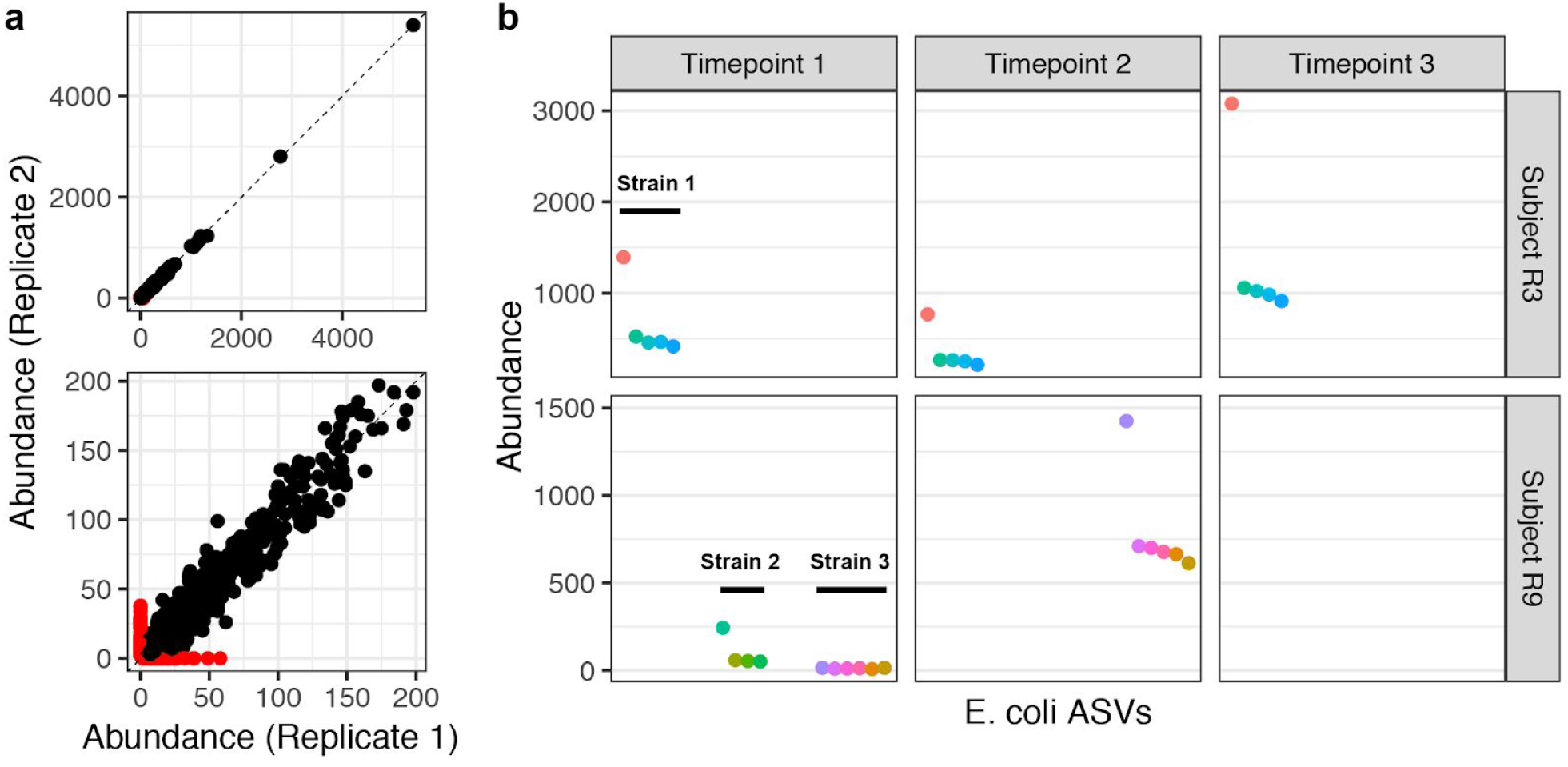
Consistent detection of full-length 16S rRNA gene sequences with single-nucleotide resolution in human fecal samples. Seven human fecal samples were characterized by two technical replicates of our full-length 16S rRNA gene amplicon sequencing method. (a) The abundances of all ASVs recovered from the same sample in different technical replicates. ASVs recovered in both technical replicates are black, and non-replicated ASVs are red. The top panel shows the full range of per-sample abundances, and the bottom panel zooms in on low abundance ASVs. (b) The abundance of each ASV assigned to *E. coli* in Replicate 2 for samples with >0.2% *E. coli* reads. Longitudinal samples from subjects R3 (top row) and R9 (bottom row) are ordered left-to-right by sampling time. ASVs putatively derived from the same strain are plotted adjacently.

To investigate whether the full complement of 16S rRNA alleles within individual strains could be resolved in more complex natural communities, we focused on all ASVs in the fecal samples that were classified as *Escherichia coli*. In each sample in which an appreciable number of *E. coli* reads was detected, clear bins of ASVs from the same strain could be constructed based on the expected integral ratios between the abundances of intra-genomic alleles, and our knowledge that *E. coli* has 7 copies of the 16S rRNA gene (Figure 5B). Consistent strain-level bins persisted over time in both subjects. In Subject R9, two different *E. coli* strains can be clearly distinguished from one another in the Timepoint 1 sample. The full complement of ASVs was recovered from the low-frequency *E. coli* strain even though the individual ASVs were present in only 8–15 reads each. The ASVs from the low-frequency strain in Timepoint 1 exactly match the ASVs from the single strain present in Timepoint 2.

## Discussion

Currently, community profiling of the 16S rRNA gene is almost always conducted using short-read sequencing technologies that measure only fragments of the complete gene. This is largely responsible for the well-known difficulty in achieving species-level resolution from high-throughput 16S sequencing data (Edgar 2018). The potential for species-level classification from full-length 16S rRNA gene amplicon sequencing has been convincingly demonstrated (e.g. Earl 2018), but higher costs, higher error rates and a less-developed ecosystem of computational methods continue to limit the appeal of long-read amplicon sequencing. We believe that recent and ongoing improvements in the accuracy and cost-efficiency of long-read sequencing technologies, coupled with the high-resolution computational methods now being developed, make a compelling case for investigators to revisit long-read amplicon sequencing as a measurement technique going forward.

The resolution and accuracy achieved by our methodology derives in significant part from the exceptional and not-entirely-appreciated accuracy of modern PacBio CCS sequencing. We observed total error rates of 4.3 × 10^−4^ per nucleotide in PacBio CCS amplicon sequencing reads, significantly *lower* than the per-base error rates of common Illumina sequencing platforms (Schirmer 2016, Pfeiffer 2018). As a result, half of all sequencing reads were error-free over the entire ~1.5 kilobase (kb) 16S rRNA gene and a computational approach leveraging repeated observations of error-free sequence was adaptable to PacBio CCS data (Callahan 2016; Callahan 2017). We predict that a computational workflow based on the DADA2 method will continue to be effective for PacBio CCS amplicons extending out to ~3 kb but that sensitivity to low-frequency variants will degrade for >3 kb amplicons as the fraction of error-free reads declines. In the regime in which few or no sequences are expected to be error-free, alternative computational methods will be necessary and are now being developed (Kumar 2019). We urge caution in applying our computational methods to data from PacBio RSII sequencing chemistries before P6-C4 and/or CCS data that was generated by the earlier SMRT Portal software, as error rates in such data may be substantially higher than in the data considered here. In addition, the 16S rRNA gene is almost devoid of long homopolymeric repeats, so additional validation is warranted if applying our methodology to more homopolymer-rich genetic loci given previous reports of higher PacBio CCS error rates in homopolymer regions (Francis 2018; Wenger 2019).

One of the challenges in evaluating our method was that the accuracy and completeness of the ASVs we recovered often outstripped supposedly authoritative references. This was particularly true when reference genome assemblies were based on short-read sequencing, which struggles to resolve repeated regions such as the multiple copies of the *rrn* operon present in many bacterial genomes, as exemplified by our re-analysis of the Wagner 2016 results. For this reason we used a multifaceted approach to evaluate ASV accuracy, in which we compared ASVs to the references provided with the mock community materials and to broader reference databases such as nt, and also investigated the pattern of integral ratios expected between genuine allelic variants within a genome. We observed very high ASV accuracy in both the Zymo and HMP mock communities (no false positives, all taxa detected) and very low residual error rates (zero – 2.6 x 10^−6^) that compare favorably to the best error rates reported from short-read amplicon sequencing (e.g. Kozich 2013; Callahan 2016). These results suggest that our methodology will be useful in applications that demand the highest levels of accuracy, such as generating reference-quality gene sequences, particularly for multi-copy genes that may have allelic variation that is difficult to resolve from short-read shotgun sequencing data. The short turn-around time of PacBio sequencing and the effectiveness of amplicon sequencing in mixed and heterogeneous samples also suggest potential diagnostic applications (Cummings 2016).

Single-nucleotide resolution of the full-length 16S rRNA gene often reveals intragenomic allelic variation. At first glance this may appear to be an unwanted nuisance, at least for simple applications such as counting the number of microbial types in a community. However, the full allelic complement can qualitatively improve the taxonomic resolution afforded by 16S rRNA gene sequencing, as we demonstrated here by identifying an *E. coli* strain as belonging to the enterohemorrhagic O157:H7 clade that is a major foodborne pathogen and threat to public health (Lim 2010). Short-read 16S rRNA gene sequencing would identify the same strain only as belonging to the *Escherichia/Shigella* genus, and traditional OTU binning approaches applied to full-length 16S rRNA gene data would group this pathogenic *E. coli* strains with other non-pathogenic *E. coli* strains in a shared OTU. New methods that automate the simple rules we used to create strain-level bins in our data by hand – integral ratios, similar taxonomic assignments, and consistent patterns across samples – will help realize the potential of full-complement classification. This could perhaps be achieved by translating methods developed for metagenomics binning (e.g. Alneberg 2014) or distribution-based clustering of marker gene sequences (Preheim 2013; Frøslev 2017). The greatest challenge to maximizing the classification potential of long-read amplicon sequencing may be the development of databases that include the full complement of 16S rRNA alleles for each reference entry.

In recent years, additional sequencing technologies have become available that could be used for long-read amplicon sequencing. Oxford Nanopore sequencing technology can achieve reads of 10s or even 100s of kilobases in length, and offers compelling benefits such as portability and rapid data acquisition. Recent work has examined Oxford Nanopore sequencing of the full-length 16S rRNA gene with promising results (Calus 2018). However, at this point in time the error rates of Oxford Nanopore remain too high to achieve single-nucleotide resolution from community samples. While new molecular consensus sequence approaches are being developed, the available approaches for Oxford Nanopore sequencing do not yet achieve anywhere near the accuracy of PacBio CCS reads. Another approach is to create “synthetic” long reads by assembling short reads known to derive from the same long amplicon molecule via some barcoding scheme (e.g. Burke 2016; Cole 2016) as recently commercialized for microbiome profiling by Loop Genomics (Wu 2019). In principle, we expect that single-nucleotide resolution and high accuracy would be achievable from synthetic long reads, as sufficient short-read coverage of long amplicon molecules can yield very low error rates. However, further validation is warranted as unexpected challenges such as systematic error modes could introduce complications.

Full-length 16S rRNA gene sequencing, short-read 16S rRNA gene sequencing, and shotgun metagenomic sequencing each have different advantages that make them preferable in different applications. Short-read amplicon sequencing is the most inexpensive taxonomic profiling method, and is also the method best suited for probing rare community variants given the great depth per-sample it can provide. Shotgun metagenomic sequencing can achieve species and even sub-species level taxonomic resolution when suitable genomic references are available, and also provides information about the functional genetic potential of the sampled community. Deep shotgun sequencing additionally allows for the de novo assembly of at least partial genomes from abundant members of the community. Full-length 16S rRNA gene sequencing combines the targeting of amplicon sequencing with species and sub-species level taxonomic resolution. We believe that amplicon sequencing of the full-length 16S rRNA gene will be an attractive option for applications that benefit from the advantages of targeted amplicon sequencing, e.g. environments in which the genetic material of interest is a minority of all DNA, and that benefit from taxonomic resolution below the genus level and/or from the breadth of available 16S rRNA gene reference databases, e.g. environments less well characterized than the human gut.

Finally, while we focused here on the 16S rRNA gene, the most important applications of targeted long-read sequencing with single-nucleotide resolution may be outside the microbial profiling application. Targeted sequencing is commonly used to characterize variation at specific loci within the large human genome, such as somatic mutations of known oncogenes in breast tumors (Chen 2018), and our method could potentially capture entire oncogenes rather than incomplete slices. Targeted sequencing of the full-length 16S rRNA gene is a standard technique to identify unknown bacteria in research and clinical settings (Woo 2008), and our method could provide the same information without the purification needed for Sanger sequencing, which is particularly useful when dealing with polymicrobial samples or unculturable bacteria (Cummings 2016). Our method may even scale to sequencing of the entire ~5 Kb 16S-ITS-23S gene region in low-complexity samples or clonal isolates. Perhaps most intriguing, multi-kilobase reads are sufficient to capture complete viral genes and even some complete viral genomes. Binned long-read amplicon sequencing is already being used to describe adaptive dynamics in the *env* gene of HIV-1 (Caskey 2017), an application that would clearly benefit from the ability to resolve the individual mutations that fuel within-host diversification. Other potential applications include MHC-typing (Karl 2017), broad-spectrum fungal profiling (Tedersoo 2018), and immune repertoire sequencing (Georgiou 2014), and we look forward to future developments in other areas that can leverage the precise long-read amplicon sequencing that new sequencing technologies and computational methods are making possible.

## Author Contributions

BJC designed the research; BJC implemented the algorithm; BJC performed the analysis; BJC wrote the paper; JW, CH and SO developed the amplicon sequencing methodology, performed the amplicon sequencing, and processed the raw sequencing data; CMT, ASG, SKM and MKD collected the human fecal samples.

## Data Availability

Reproducible R markdown documents implementing the analyses presented in this manuscript are available at https://github.com/benjjneb/LRASManuscript. Sequencing data are deposited in the SRA under Bioproject accession PRJNA521754 and will be made public upon acceptance.

## Acknowledgements

We thank Naga Betrapally for assistance in generating CCS reads, Meredith Ashby for advice and consultation, and Paul Hess, Roey Angel and Jacob Price for sharing test datasets not included in this manuscript. CoreBiome, Inc. performed the fecal sample DNA extraction and quantification.

## Supplementary Tables and Figures

**Supplementary Figure 1.**
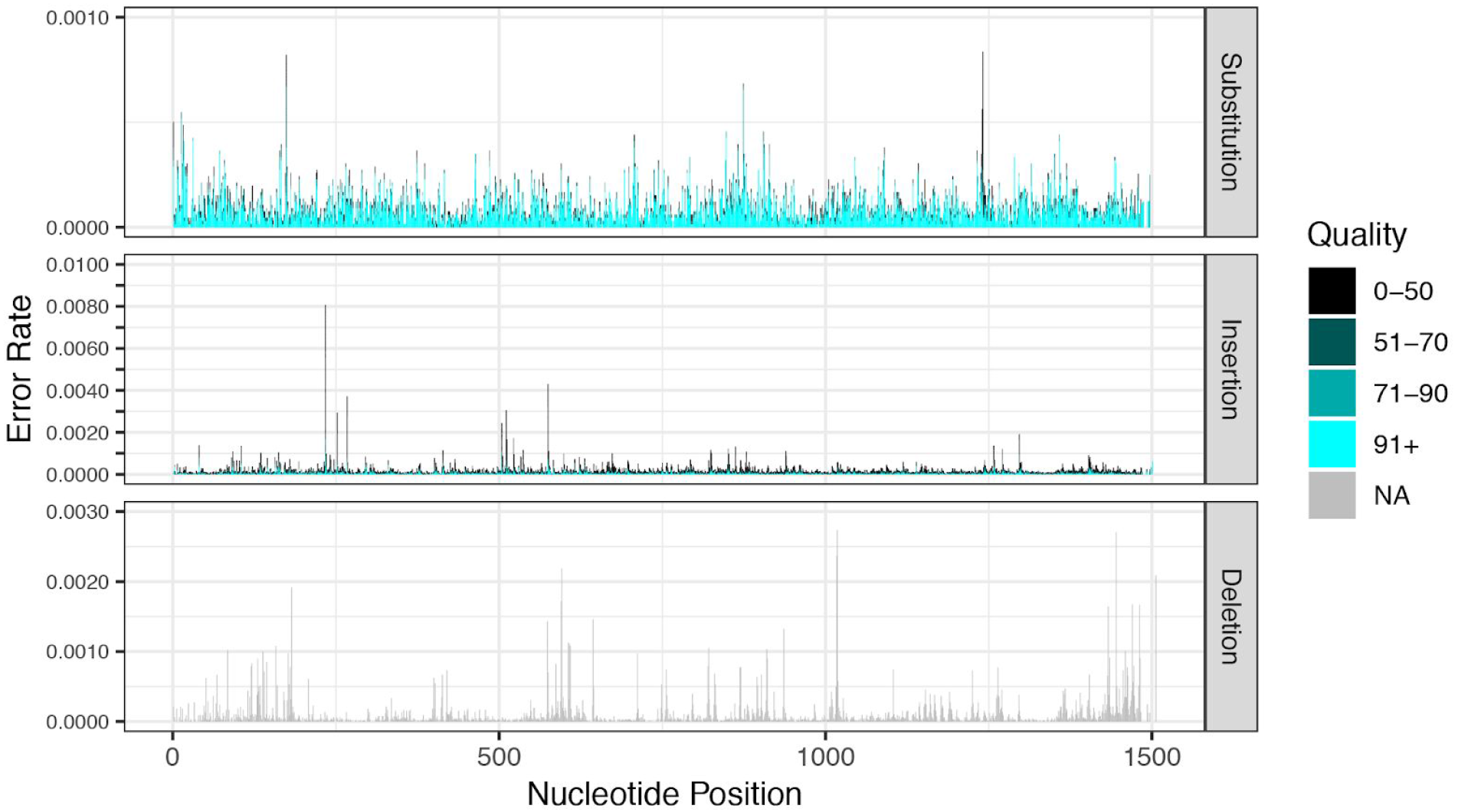
Error rates in PacBio CCS amplicon reads from the Zymo mock community as a function of error type, nucleotide position and quality score. The rate of substitutions (top), insertions (middle), and deletions across all non-chimeric and non-contaminant reads from the Zymo mock community are shown. Lower quality bases are plotted in darker colors. There is no quality score associated with deletions, as such errors indicate the absence of the corresponding nucleotide in the sequencing read.

**Supplementary Table 1.**
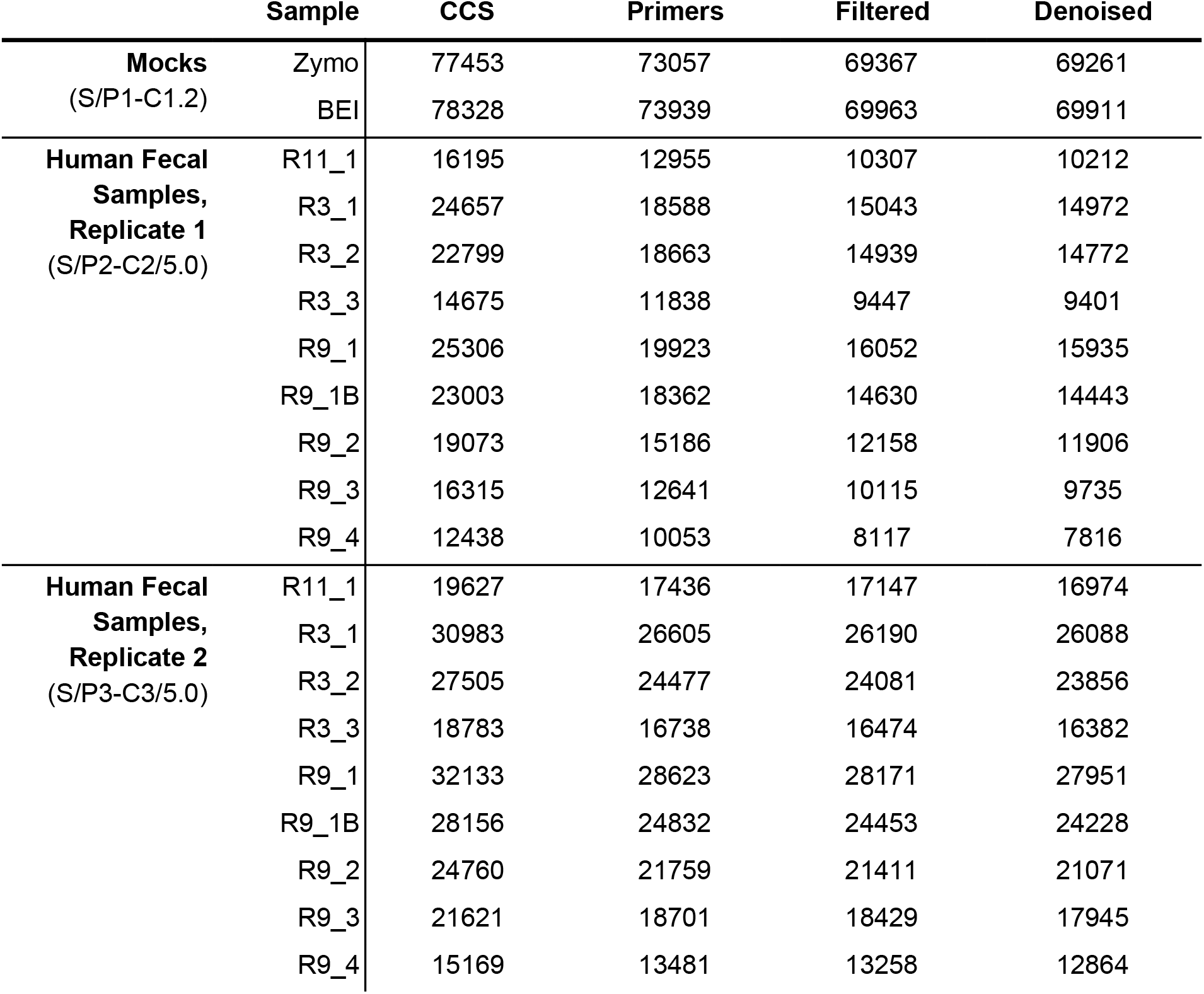
Reads retained at each step of our computational processing pipeline. Three categories of samples were included in this manuscript: mock community samples, and the first and second replicate samples derived from the human fecal specimens. The mock community samples were sequenced on dedicated Sequel cells, while the human fecal samples were 12-fold multiplexed. The sequencing chemistry used for each set of samples is indicated in parentheses. CCS: PacBio CCS reads that met the default *minPasses*=*3* and *minPredictedAccuracy*=*0.999* thresholds for inclusion. Primers: PacBio CCS reads in which the forward and reverse primer sequences were detected with at most two mismatches each. Filtered: PacBio CCS with primers that passed the *maxEE*=*2*, *minLen*=*1000* and *maxLen*=*1600* filtering thresholds. Denoised: PacBio CCS reads that were successfully denoised.

|  | Sample | CCS | Primers | Filtered | Denoised |
| --- | --- | --- | --- | --- | --- |
| **Mocks**<br>(S/P1-C1.2) | Zymo | 77453 | 73057 | 69367 | 69261 |
|  | BEI | 78328 | 73939 | 69963 | 69911 |
| **Human Fecal Samples, Replicate 1**<br>(S/P2-C2/5.0) | R11_1 | 16195 | 12955 | 10307 | 10212 |
|  | R3_1 | 24657 | 18588 | 15043 | 14972 |
|  | R3_2 | 22799 | 18663 | 14939 | 14772 |
|  | R3_3 | 14675 | 11838 | 9447 | 9401 |
|  | R9_1 | 25306 | 19923 | 16052 | 15935 |
|  | R9_1B | 23003 | 18362 | 14630 | 14443 |
|  | R9_2 | 19073 | 15186 | 12158 | 11906 |
|  | R9_3 | 16315 | 12641 | 10115 | 9735 |
|  | R9_4 | 12438 | 10053 | 8117 | 7816 |
| **Human Fecal Samples, Replicate 2**<br>(S/P3-C3/5.0) | R11_1 | 19627 | 17436 | 17147 | 16974 |
|  | R3_1 | 30983 | 26605 | 26190 | 26088 |
|  | R3_2 | 27505 | 24477 | 24081 | 23856 |
|  | R3_3 | 18783 | 16738 | 16474 | 16382 |
|  | R9_1 | 32133 | 28623 | 28171 | 27951 |
|  | R9_1B | 28156 | 24832 | 24453 | 24228 |
|  | R9_2 | 24760 | 21759 | 21411 | 21071 |
|  | R9_3 | 21621 | 18701 | 18429 | 17945 |
|  | R9_4 | 15169 | 13481 | 13258 | 12864 |

